# A Zymography technique to study amino acid activation by aminoacyl tRNA synthetases (aaRS): A broad spectrum, high-throughput tool to screen activities of aaRS and their “Urzyme” variants

**DOI:** 10.1101/2023.02.01.526722

**Authors:** Sourav Kumar Patra, Charles W. Carter

## Abstract

Amino acyl tRNA synthetases or aaRSs play a key role in assuring the precision of protein translation. They are highly specific for their cognate amino acid and cognate tRNA substrates during protein synthesis, utilizing ATP to ensure that proper assignments are made between amino acid and anticodon. Specific aaRS for each amino acid are present in all cells. We describe a new zymography technique to qualitatively visualize and semi-quantitatively determine the amino acid activation capacity of each type of aaRS molecule by indirect colorimetric detection of released pyrophosphates during the formation of aminoacyl-AMP. Protein samples containing aaRS are subjected to Native PAGE, followed by incubation in buffer containing cognate amino acid and ATP for sufficient time to generate pyrophosphates (PPi) which are then converted to inorganic phosphates by pyrophosphatase treatment. Finally, the generated and localized phosphates around the aaRS protein inside the gel can be visualized after staining by ammonium molybdate and malachite green solution. This technique has been validated by inspecting the substrate specificities of specific aaRSs. This zymography technique is sufficiently sensitive to detect and authenticate activities of much (i.e., ~10^-5^-fold) less active aaRS “Urzymes”, to study alteration of activities of aaRS by various intrinsic or extrinsic factors and to screen aaRS-specific antimicrobial drugs.

## 1. Introduction

Zymography is considered the most authentic way to visualize[1] the localized activity of an enzyme or its isoforms[2] present in a mixture of proteins (cell lysate or semi purified protein sample) using polyacrylamide gel electrophoresis. Successful zymography requires only a proper interface or step for enzyme substrate interaction and a colorimetric detection method of either product formation or loss of substrate by a proper choice of staining reagent. In the past few decades several established zymography methods for different proteases[3, 4], lipases[5], cell wall degrading enzymes[6–9], redox active enzymes[10] etc. played key roles in biological research.

Aminoacyl tRNA Synthetases (aaRS) are an ancient group of 20 proteins, specific for each amino acid. Their crucial role in protein translation[11–13] is to charge the corresponding amino acid with ATP (amino acid activation) and transfer it to the terminal 2’ or 3’ ribose - OH group of cognate tRNA molecules (aminoacylation), thereby completing the assignment of amino acid to anticodon required by the universal genetic code. Due to their high specificity towards substrates (cognate amino acid and tRNA), aaRSs are major contributors to the precision of protein translation. For that reason, they are also important antimicrobial drug targets[14]. Despite its potential value, no zymography method has been demonstrated for aaRS. The technique described here is useful to visualize and study catalysis of amino acid activation; the first step of the consecutive two step reactions executed by aaRSs.

In the amino acid activation reaction, an aaRS utilises ATP to activate its cognate amino acid through formation of [aminoacyl-5’AMP] and release of PPi. This released PPi is enzymatically converted to orthophosphates (PO_4_^-^), which are then visualised by the staining method described here to visualize the amino acid activation reaction by that specific aaRS.

In this newly developed zymography technique using Native PAGE, amino acid activation by specific aaRSs is demonstrated for the first time using both purified aaRS and crude cell lysates. The authenticity of this technique is also established by demonstrating amino acid– and ATP–dependence of different aaRSs. The simple execution, high sensitivity, and short time frame required for zymography make this technique unique and useful.

Ammonium molybdate has long been used to detect phosphates in analytical biochemistry. In this new method, Malachite green and hexaammonium heptamolybdate tetrahydrate greatly enhance the intensity of the coloured complex with orthophosphates generated from PPi above and adjacent to active aaRS proteins in the gel[15]. The Malachite green assay was previously used by several groups to study cyclic nucleotide phosphodiesterase activity[16], phosphoserine phosphatase activity[17], library screenings for kinases[18] and, especially to measure primitive aminoacyl-tRNA synthetase (protozyme) activity[19]. We describe here the first adaptation of this assay into a simple, rapid, and sensitive zymography technique, with specific protocols, cognate amino acid-dependence, and insensitivity to phosphatase and kinase blockers, so that aaRS activities can be detected in both complex protein mixtures and purified samples.

## 2. Materials and methods

### 2.1. Strains and constructs used

Purified full-length wild type Leucyl-tRNA synthetase (LeuRS), Tryptophanyl-tRNA synthetase (TrpRS) and induced and overexpressed full-length Histidyl-tRNA synthetase (HisRS) were used. Maltose binding protein (MBP) fused LeuAC and HisRS-4 “urzymes” were also used. MBP-LeuAC urzyme was kept in pMAL-c2X plasmid and transformed into BL21(DE3)pLysS competent cells before overexpression and purification. HisRS urzyme construct was also overexpressed using BL21(DE3)pLysS. Purification of the proteins was done as per Hobson et. al[20], Li et al[21], Williams et al[22].

### 2.2. Cell lysis buffer preparation and Cell Lysis

Cell lysis buffer, 20 mM Tris-HCl of pH 7.5 was diluted from a stock of 100 mM Tris-HCl at pH 7.5, to which EDTA and Lysozyme were added to the final concentration of 1 mM and 1 mg/ml respectively. 1 tablet of pierce protease inhibitor cocktail was also added for making 50 ml of Cell lysis buffer. Bacterial Cells were lysed, by sonication with 8 pulses of 10 second 70% amplitude sonic vibrations keeping 20 seconds of pause time, ensuring the tube remained in ice during sonication. Lysed cells were then centrifuged at 12000 rpm for 10 min at 4 °C and supernatants were collected, aliquoted in tubes, and stored at −80°C [20, 23–25].

### 2.3. Sample Loading dye preparation

A 5X sample loading dye contains 62.5 mM Tris-HCl of pH 6.8; 40% (v/v) glycerol and 0.01% (w/v) Bromophenol Blue. Commonly used reagents such as 2-Mercaptoethanol (β-mercaptoethanol) or any reducing agents as well as SDS or any other detergent should be avoided as they may destabilize structures, inhibiting activity.

### 2.4. Native gel Electrophoresis buffer preparation

For making 1 litre of electrophoresis buffer, weigh 3.03 grams of Tris and 14.4 g of glycine, then mix it in deionized H_2_O and make the volume up to 1 litre in a graduated cylinder. This must be made fresh on the day of experiment or just before the day and should be kept at 4 °C. Do not add SDS.

### 2.5. Preparation of Native gel

Unlike the standard SDS PAGE, for this method, 8% (v/v) resolving and 5% (v/v) stacking gel should be made without any addition of SDS[2].

### 2.6. Substrate reaction buffer preparation

Make a substrate reaction buffer of 50 mM HEPES of pH 7.5, 5 mM ATP, 100 mM cognate amino acid, 20 mM MgCl_2_ and 50 mM KCl. For cell lysates or complex protein mixtures, add phosphatase and kinase inhibitor cocktail (SIGMA-Aldrich) in the substrate reaction buffer mix. When screening a drug or any other compound’s effect on aaRS activity, the substrate reaction buffer can also include that compound to check the relative activities of aaRS in a single or multiple incubation setup. Moreover, the pH of the incubation buffer, the amount of ATP and the type or amount of amino acid may be different, depending on the user’s need and experimental goals.

### 2.7. Preparing the malachite green-ammonium molybdate staining solution

Separately make 0.05% (w/v) Malachite green in 0.1 N HCl and 5% (w/v) hexaammonium heptamolybdate tetrahydrate solution in 4N HCl first. Then Mix Malachite green and ammonium molybdate solution in 3:1 ratio and mix well by stirring for 20 minutes. After that filter the solution and store it at 4°C inside a clean, closed, dark glass container.

### 2.8. Procedure of Zymography

#### 2.8.1. Sample preparation

Sample preparation for this technique should be done freshly the day of experiment so as to retain the activity of enzymes as well as their conformations intact. Mix the protein sample with native sample loading dye and adjust the sample volume with native gel running buffer inside a 0.5 ml microcentrifuge tube. Sample preparation must be done without heat treatment and the sample(s) should be kept on ice to keep enzyme activity intact until subjected to electrophoresis. If testing a cell lysate, make samples using at least 20 μg or more protein for each lane for optimal results. Lower protein content (2-3 μg of protein) suffices for purified or semi purified proteins. For quantitative as well as qualitative analysis, always normalize the protein content of samples before subjecting to electrophoresis. Protein concentrations should be measured using standard protein concentration measurement kits preferably using Bradford reagent of Bio-RAD using a protein standard before sample preparation.

#### 2.8.2. Sample loading and Electrophoresis process

Before sample loading and electrophoresis, always pre-electrophorese for 30 minutes at 40 mA keeping the gel setup at 4 °C. This step will remove residual TEMED, APS and products of incomplete polymerization that might interfere with mobility of proteins through the gel and also with their activity. Then load and run the samples in the native gel running buffer or electrophoresis buffer at constant current flow of 30-40 mA keeping the system at 4 °C. once the dye font reaches at the bottom of the gel, continue the electrophoresis process for 30 minutes to 1 hour more This step ensures the complete entry of desired protein and thus needs to be standardised according to the user’s desired aaRS and its migration through the gel. A parallel gel is useful for Coomassie blue staining.

#### 2.8.3. Enzyme-substrate incubation step

After electrophoresis, remove the gel carefully and rinse it with deionized autoclaved water while keeping it in a clean small glass box on a gyratory or orbital shaker for 5 mins. This rinsing step can be repeated twice. This step will remove the electrophoresis buffer, glycine and other unwanted particles from gel. After washing the gel with deionized water, pour the substrate reaction buffer into the glass box containing the gel and keep the setup on a shaker for 45 minutes at 4°C. This step will ensure the soaking of substrate mixture inside the gel; the low temperature will minimize the enzyme activity and helps prevent diffusion of product (pyrophosphate) into the other parts of gel and solution, thus increasing the contrast after staining of gel. After 45 minutes add polyethylene glycol (PEG-8000; Sigma-Aldrich, Cat. No. 25322-68-3) into the mixture to a final concentration of 5% −8% (w/v) and shake for another 15 minutes until the PEG totally dissolves and is absorbed by the gel. Polyethylene glycol will reduce diffusion, helping to localize pyrophosphates and inorganic phosphates within the gel.

Discard a maximum amount of substrate buffer mix from the gel containing glass box and transfer the gel-box inside a 37°C incubator covering with cellophane paper or Saran wrap to minimize evaporation and drying. Make sure that the gel is submerged but not covered up by substrate buffer mix. Try to keep minimum amount of substrate buffer mix to avoid drying and shrinking of gel or if possible don’t keep any amount of buffer mix at all if incubating for short times. Keep the gel like this at 37°C for at least 1 hour to check wild type full length aaRS enzyme activity. To detect urzymes or any other low activity having aaRS variant incubate the gel for at least 2 hours at this condition.

Then carefully add pyrophosphatase solution [NEB] (0.1Unit/ml) on the gel dropwise by pipetting, keeping in mind that the whole gel should be covered. Keep the gel in this condition for 30 minutes more at 37°C. pyrophosphatases will convert most of generated pyrophosphates during aminoacyl-AMP formation into phosphates.

#### 2.8.4. Staining process

After the incubation step, decant all residual substrate buffer mix or pyrophosphate solution and add an adequate amount of staining solution of hexaammonium heptamolybdate tetrahydrate and malachite green mixture just to cover the gel. Shake the gel box gently for 15 minutes at room temp. The gradual development of dark greenish bands around aaRS protein present in the gel will signify the phosphomolybdate-malachite green complex formation and thus the activity of aaRS protein(s). Capture the image of the gel using designated scanner (for taking colour picture) or use Gel-doc imager (BioRAD) for taking greyscale image. The complex of phosphomolybdate-malachite green has an absorbance maximum at 620 nm - 650 nm range of light. So, use of more advance imaging system having customized wavelength setup can also increase the contrast of imaging and highlight the band(s). Imaging of the gel should be done in highly acidic staining solution. Due caution should be used to avoid exposure to the highly acidic solution during these steps of gel staining, band development and imaging.

#### 2.8.5. Semi-quantitative or qualitative Analysis

For semi-quantitative or qualitative analysis, a chromatic band intensity can be measured using ImageJ software [25] to determine relative activities of aaRS molecules in different samples.

## 3. Results

### 3.1. Malachite Green-ammonium molybdate staining detects induction of full length aaRS as well as much less active aaRS urzymes in crude cell extracts

The malachite green assay used by Onodera et al.[19] in studying the amino acid activation by 46-residue HisRS and TrpRS protozymes illuminated the pathway for adapting, improvising and developing a new zymography technique to study the aaRS amino acid activation capacity *in situ* after separation on a native PAGE gel. Purified wild type aaRS proteins showed significant band formations. However, a greater challenge lay in identifying specific aaRS activities in complex mixtures of proteins. We demonstrated full-length wild type *E. coli* HisRS activity in both IPTG uninduced and induced cell lysates **(Fig. 1)**.

**Fig. 1.**
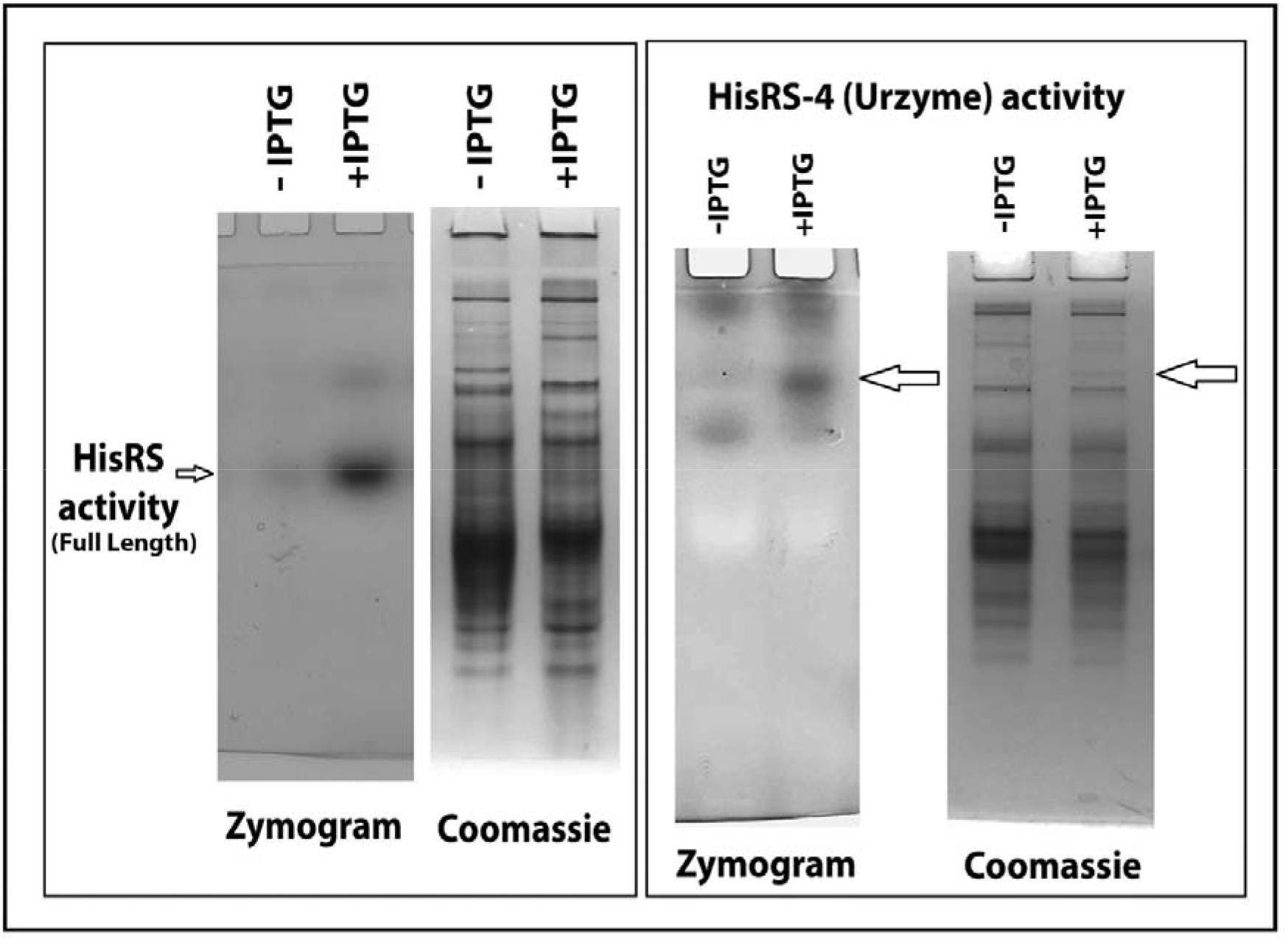
Detection of full length Histidyl tRNA synthetase (HisRS) activity and less active HisRS-4 urzyme activity in crude cell lysates confirm and authenticate the specificity of aaRS zymography technique as well as its sensitivity. Cell lysates of IPTG uninduced and induced BL21(DE3)pLysS competent cells transformed with *E. coli* HisRS (full length Histidyl-tRNA synthetase) and MBP-HisRS-4 “Urzyme” was used separately during this study. 15 μg of total protein was loaded in each well of an 8% native gel from IPTG treated and untreated cell lysates. Another gel was electrophoresed in the same way and used for Coomassie staining.

### 3.2. Malachite Green-ammonium molybdate zymography has high sensitivity

Urzymes (“ur” meaning primitive, original, earliest) constructed from both aaRS Classes contain only the amino acid activation and acyl-transfer active sites of the full-length enzymes. However, their activity is reduced by up to 5 orders of magnitude, relative to the full-length enzymes from which they were derived. Nevertheless, an analogous comparison of uninduced and IPTG induced MBP-linked HisRS-4 urzyme showed significantly enhanced activity in the induced sample, highlighting the high sensitivity of this zymography technique as reported by Onodera, *et al*. (**Fig. 1**). The presence of multiple activity bands in these gels suggests the existence of multimeric forms of both full length aaRSs and urzymes.

### 3.3. AARS zymography is both amino acid and ATP-dependent

Specific aaRS activities can be visualized using cognate amino acids in substrate incubation buffers (**Fig. 2**). Activity staining of wild type full length aaRS is much less time consuming and more prominent in this method because of their high activity in gels. We authenticated the aaRS activity from the point of view of substrate specificity by using two different types of full-length wild type purified aaRS proteins, TrpRS and LeuRS. This was done using leucine and tryptophan as cognate amino acid substrates alternatively in two different incubation setups. LeuRS showed activity in the gel setup where leucine was used but not tryptophan whereas, conversely TrpRS showed high activity only when incubated tryptophan, as expected. ATP is the other substrate for amino acid activation by aaRS. We used MBP-LeuAC “urzyme” to demonstrate the ATP-dependence of aaRS zymography, reinforcing the authenticity of the activities of only aaRS but not any other proteins (**Fig.3**).

**Fig. 2.**
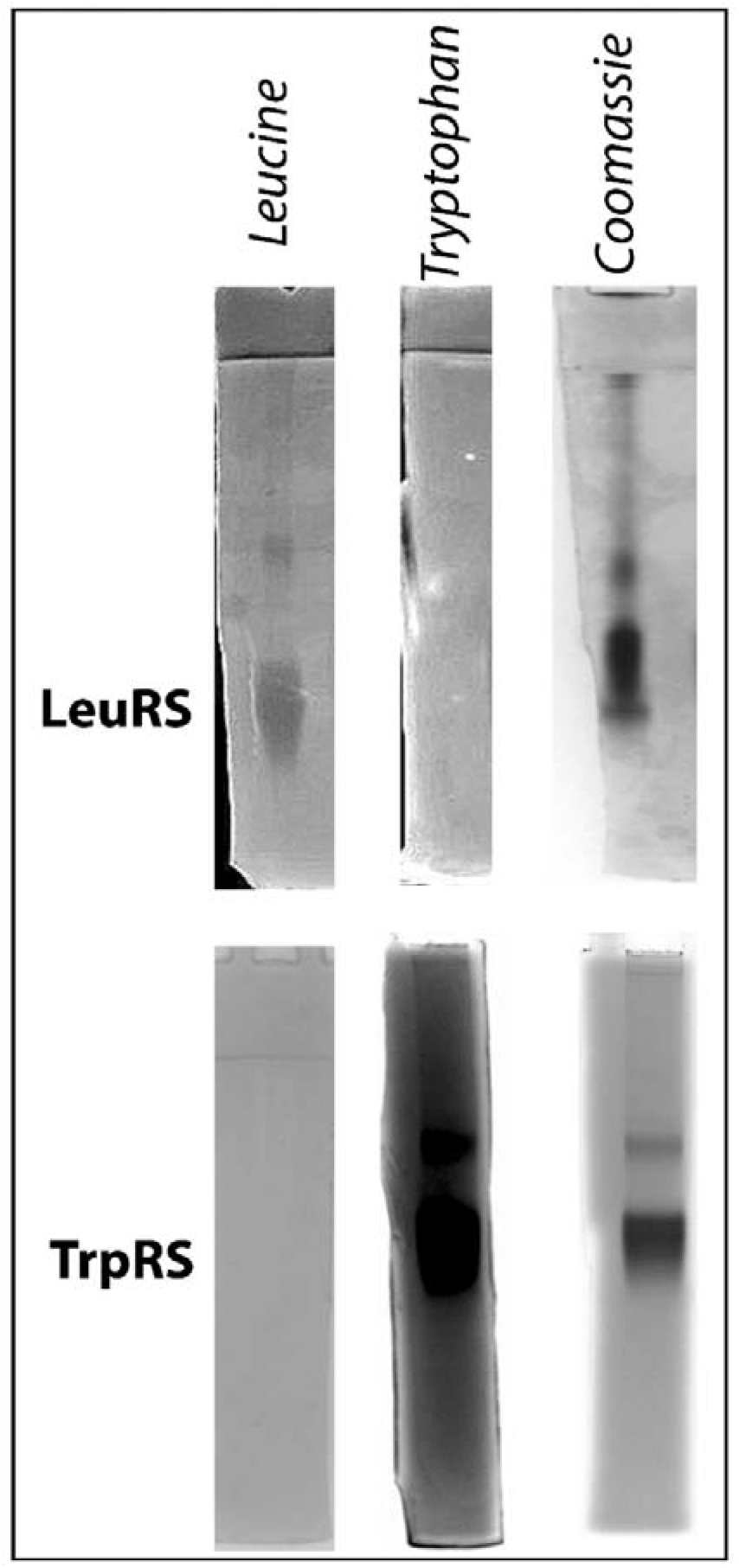
Different full length aaRSs (LeuRS and TrpRS) and alternative cognate amino acids (Leucine and Tryptophan authenticate aaRS zymography and substrate specificity. To demonstrate substrate specificity, purified LeuRS and TrpRS were used in native PAGE. After electrophoresis process, different cognate amino acids were used alternately during gel incubation in different setups followed by pyrophosphatase treatment and staining of gels. 2 μM of each enzyme were used for this experiment. A parallel gel was run with same amount of protein in the same setup for Coomassie staining.

**Fig. 3.**
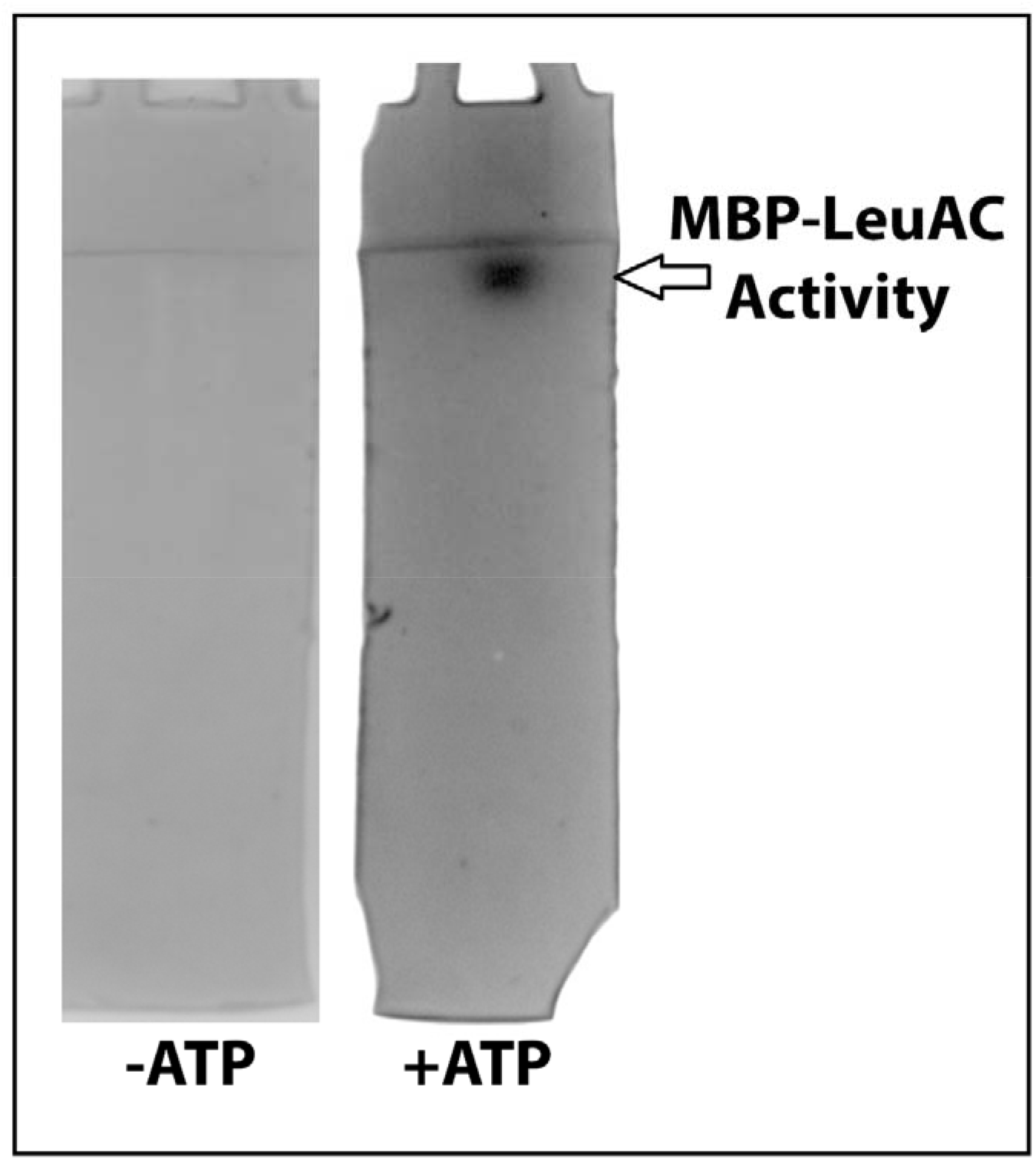
Authentication of aaRS zymography by showing the activity dependence on ATP, the second substrate for aaRS enzymes. MBP-LeuAC enzyme was used for this ATP dependence study. 10 μM of MBP-LeuAC was loaded in each well and the electrophoresis was done for 1 hour at 40mA. After that, the gels were incubated in incubation buffer having everything (100 mM leucine & 5 mM ATP) in one set and in another set, with everything except ATP. Subsequent pyrophosphatase treatment and staining showed a sharp, achromatic band of MBP-LeuAC activity in the setup incubated with, but not without ATP, confirming the ATP dependence of this reaction and thus the authenticity of this technique.

## 4. Discussion

This zymography method is a straightforward way to confirm amino acid activation by a specific aaRS enzyme by showing that staining is dependent on provision of specific cognate amino acid and ATP substrates. It can be useful to identify relative expressions, hence activity of a particular aaRS, by staining different biological samples or treatments in different lanes in the same Native PAGE by normalizing the total protein in each lane.

A decisive benefit of the sensitivity of this technique entails authenticating the catalytic activity of engineered variants. A foremost example is aaRS urzymes, which are smaller, much less active, minimal catalytic domains excerpted from full length aaRS enzymes proposed to represent primitive ancestral aaRS forms[20, 21, 26]. Though they contain scarcely more than a hundred amino acids, urzymes retain far smaller magnitudes of the basic catalytic capabilities of the synthetase from which they were derived[27]. We will address this key result in greater detail in a separate publication.

Other significant applications include activity screening of recombinant or engineered aaRS proteins for various purposes, including the use of protein design to screen for more active, more soluble, or more stable aaRS urzymes in crude extracts. The ability to screen combinatorial libraries in crude extracts will potentially enable higher throughput processing of such design efforts.

Potential antimicrobial drugs targeting aaRS enzymes could also be screened by this compact technique. If proper incubation conditions can be devised, the technique is even effective to screen the effect of temperature, pH or other external agents on aaRS activity.

Finally, this technique may also be useful to reveal alterations of activity (if any) due to post-translational modifications of aaRS proteins[23, 25]. These post translational alterations might be related to oxidative[28] or nitrosative stress[29] or other stable biochemical modifications that occur *in-vivo* or *in-vitro* provided these alternations are not affected or reversed during zymography procedure of prestaining steps.

## 5. Conclusions

The straightforward execution of this technique and validation of it using purified and complex mixture of proteins, different cognate amino acids, presence and absence of ATP and the highly-sensitive detection of much less reactive aaRS urzymes strongly indicates the development of a technique which has a broad-spectrum usage.

## Acknowledgement

We acknowledge constructive comments by Guo Qing Tang.

## Author contributions

SKP conceived the work and performed the experiments described. CWC Jr suggested using PEG to limit diffusion during the staining. SKP wrote the initial draft and all authors contributed to the final draft.

## Graphical Abstract

### Schematic representation of the zymography technique of amino acid activation by aaRS

Specific amino acid activation by aminoacyl tRNA synthetase (aaRS) can be visualized using this straightforward, simple and rapid zymography technique which requires no special emphasis. Crude cell lysates containing aaRS or purified specific aaRS proteins or their much less active urzyme variants are electrophoresed in a native gel setup. The native gel is then incubated with buffer containing ATP and specific cognate amino acid (as per user’s need) followed by pyrophosphatase treatment and finally staining with malachite green-ammonium molybdate staining solution. Use of polyethylene glycol (PEG-8000) and phosphatase and kinase inhibitors (for cell lysate) during incubation increases the contrast of band formation and minimizes unnecessary bands.

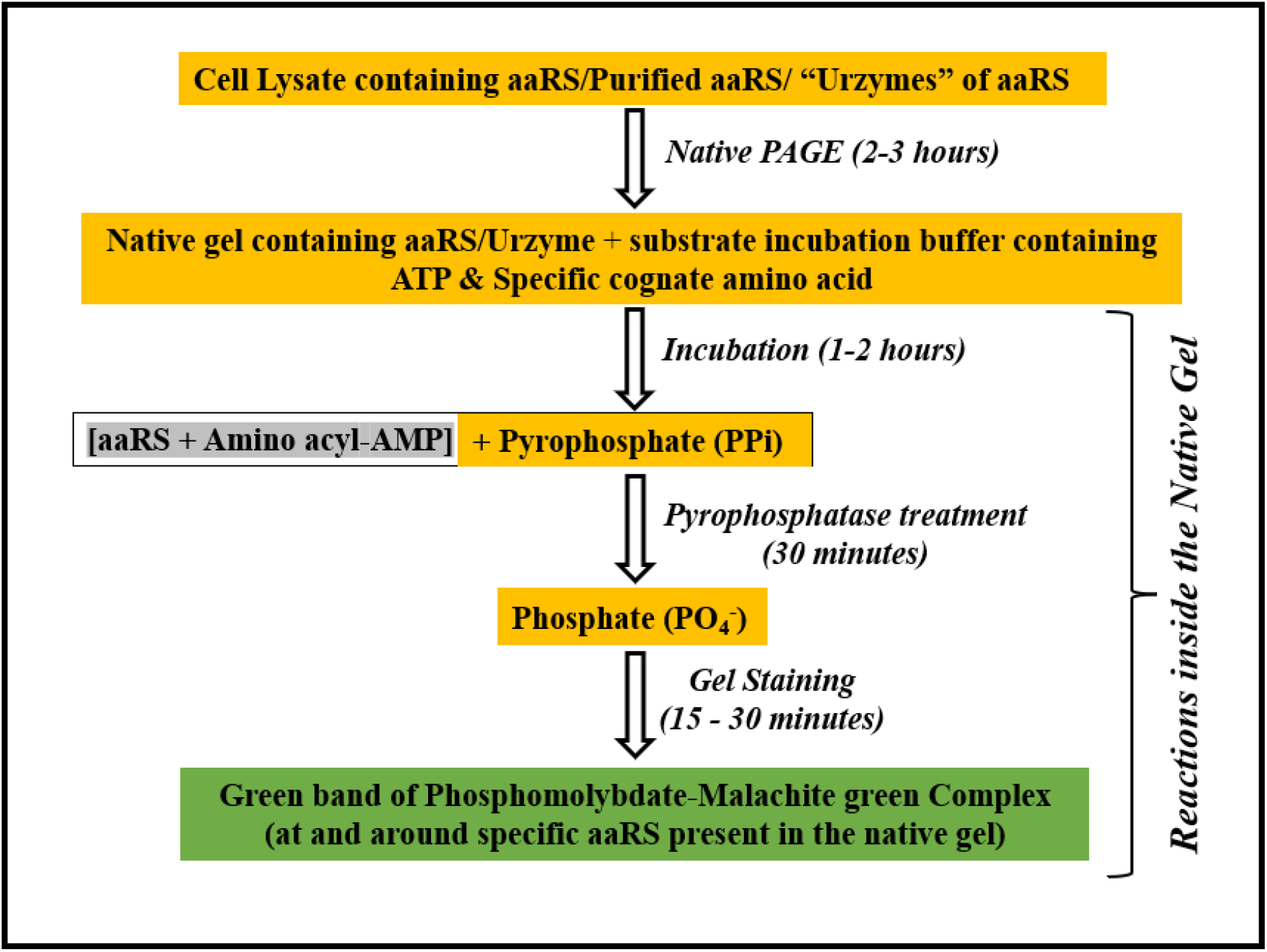

### Schematic representation of the zymography technique of amino acid activation by aaRS

Specific amino acid activation by aminoacyl tRNA synthetase (aaRS) can be visualized using this straightforward, simple and rapid zymography technique which requires no special emphasis. Crude cell lysates containing aaRS or purified specific aaRS proteins or their much less active urzyme variants are electrophoresed in a native gel setup. The native gel is then incubated with buffer containing ATP and specific cognate amino acid (as per user’s need) followed by pyrophosphatase treatment and finally staining with malachite green-ammonium molybdate staining solution. Use of polyethylene glycol (PEG-8000) and phosphatase and kinase inhibitors (for cell lysate) during incubation increases the contrast of band formation and minimizes unnecessary bands.

